# Interval timing clock property in the rat granular retrosplenial cortex

**DOI:** 10.1101/2024.06.17.598602

**Authors:** Tohru Kurotani, Ken’ichi Nixima, Tomohiro Tanaka, Yoshio Sakurai, Kazuo Okanoya

## Abstract

The rodent granular retrosplenial cortex (gRSC), densely interconnected with the hippocampal formation and the anterior thalamic nuclei, plays an important role in learning and memory. We had revealed that small pyramidal neurons in the superficial layers of the rat gRSC exhibit late-spiking (LS) firing properties. It has been suggested that neural circuits containing LS neurons can encode time intervals on the order of seconds, known as “interval timing”. To test the possibility that the rat gRSC is involved in the processing of interval timing, we employed a trace fear conditioning paradigm in which the conditioned stimulus (CS) and the unconditioned stimulus (US) were temporally separated. First, we examined the effect of cytotoxic lesions made in the RSC prior to trace fear conditioning. We found that intact rats exhibited freezing behavior after CS tone presentation, whereas lesioned rats did not exhibit such freezing behavior. Next, we conducted *in vivo* chronic or acute recordings of neural activity from the rat gRSC in a test session conducted one week after the conditioning. In both recordings, we observed a distinct spike activity in which there was a transient increase in the firing rate around the presentation of the CS tone, followed by a rapid suppression and then ramping activity (a gradual elevation of the firing rate) until the next CS presentation. This “ramping activity” is thought to be one way in which interval timing is represented in the brain. Post stimulus histogram analysis revealed the existence of ramping activity in the gRSC, which reached its peak at various time intervals after the onset of the CS tone. Interestingly, this activity was specifically observed in response to the CS tone but not to the non-CS tone. Moreover, in naive rat gRSC (no trace fear conditioning), no such ramping activity was observed. These results indicate that gRSC neurons can encode time information on the order of tens to hundreds of seconds, integrating incoming sensory input with past memory traces.

## 1. Introduction

The perception of the “how much time has passed” and the memory of the “order in which events occurred” play a crucial role in the development of survival strategies in organisms. They can use this information to optimize their behavior and adapt to their environment (Gibbon and Church, 1981, Gibbon et al., 1984, Clayton and Dickinson, 1988, Henderson et al., 2006, Ferkin et al., 2008). However, there are no sensory receptors or sensory areas of the brain that are specialized for time perception; various areas of the brain have been reported to engage in different ways to encode, memorize and refer to time information (Buhusi and Meck, 2005, Paton and Buonomano, 2018, Patke et al., 2020). Three mechanisms for encoding time in organisms have been proposed, each handling a different time scale: the sub-second order, the second to minute/hour order and the circadian rhythm (Buhusi and Meck, 2005).

The most well-studied of these, the circadian rhythm, has been elucidated as a series of mechanisms by which the information of the daily light/dark cycle is transmitted from the retina to the neural network centered in the suprachiasmatic nucleus. Activity in the network controls the endocrine system and molecular mechanisms, including gene expression (Patke et al., 2020, Pilorz et al., 2020).

The sub-second sense of time is involved, for example, in speech, the generation of motor patterns, and timing behavior (Thier et al., 2000, Edwards et al., 2002, Schirmer, 2004). The cerebellum, basal ganglia and cerebral cortex has been suggested to contribute to this type of time sensation by utilizing neural mechanisms such as adaptation, synchronization and entrainment of neural activity, plastic changes in synaptic transmission, and learning by a network of a small number of neurons (Paton and Buonomano, 2018).

Cognitive mechanisms on the order of seconds, minutes and hours are called “interval timing”. Similar to the other time scales described above, multiple neural representations for interval timing are widely scattered in the brain (Komura et al., 2001, Mita et al., 2009, Kraus et al., 2013, Tallota and Doyèrea, 2020). One such representation is recognized as “ramping activity,” in which the firing frequency of a specific neuron gradually increases as inputs to the neuron are time-integrated. By setting a certain threshold for this ramping activity to trigger a specific response, it is thought that the organism can measure a specific length of time. Indeed, ramping activity has been recorded in a variety of neural systems in animals performing tasks requiring interval timing discrimination. These include CA1 neurons of the rabbit hippocampus during trace fear conditioning (McEchron et al., 2003), thalamic neurons of the rat while performing a delayed stimulus reward association task (Komura et al., 2001), and neurons in the medial frontal cortex and the dorsomedial striatum neurons of the rat while performing a fixed-interval timing task (Emmons et al., 2017). In the present study, we conducted a trace fear conditioning paradigm using rats and found that interval timing was represented as ramping activity in the retrosplenial cortex (RSC).

The RSC is one of the largest cortical regions in the rodent posterior cingulate cortex, forming dense reciprocal connections with the anterior thalamic nuclei (ATN) and hippocampal formation (HCF). These three areas have been assumed to operate conjointly to support spatial learning and memory (Aggleton et al. 2010; Chen et al. 1994; Cooper and Mizumori 2001; Dumont et al. 2010; Garden et al. 2009; Pothuizen et al. 2008; Vann et al. 2009; Vogt and Miller 1983), the default mode network (Buckner et al. 2008; Uddin et al. 2009; Vann et al. 2009), and emotional evaluation of behavioral contexts (Maddock, 1999). Connections between the RSC and ATN strongly affect associative learning (Gabriel et al. 1980a, 1980b). Furthermore, the hippocampal-retrosplenial network is responsible for spatial representation (Honda et al. 2011). These findings strongly suggest that the RSC serves as a bridge, which links the ATN and HCF and integrates these two input signals. The RSC also has strong interconnections with cortical sensory areas and the anterior cingulate cortex, suggesting their role as a hub for integrating sensory and memory signal processing (Wang et al., 2016, Chrastil, 2018).

The rodent RSC is divided into two sub-regions: the dorsal dysgranular RSC (dRSC) and the ventral granular RSC (gRSC). The superficial layers of the gRSC consist of a patchy, dense modular structure of small pyramidal neurons that project to the contralateral gRSC (Ichinohe et al. 2008; Sripanidkulchai and Wyss 1987; Wyss et al. 1990). Their apical dendrites form prominent bundles that spread to layer 1a and co-localize with patchy thalamic terminations mainly from the anterior thalamic nucleus (Ichinohe et al. 2003; Odagiri et al. 2011; Shibata 1993; Sripanidkulchai et al. 1986; Van Groen et al. 1990, 2003; Wyss et al. 1990).

We have shown that these small pyramidal neurons in the superficial layers of the gRSC are “late-spiking (LS)” neurons with delayed firing properties, generating action potentials with a time delay in response to inputs (Kurotani et al., 2013). Due to this firing property, LS neurons alone may function as a time integrator on the order of seconds. Furthermore, by connecting LS neurons to each other, it may even be possible to encode a time interval longer than several tens of seconds. Thus, we hypothesized that the gRSC may demonstrate the ability to encode interval timing information.

To test this hypothesis, we conducted behavioral experiments and multi-unit neural recordings from the gRSC in rats subjected to a trace fear conditioning paradigm in which the conditioned stimulus (CS) tone and the unconditioned stimulus (US) foot shock were temporally separated by a trace interval. Rats subjected to trace fear conditioning began to show freezing responses, which started just after termination of the CS tone. This indicates that rats could learn causal relationships between CS and US even with a time interval of more than 10 seconds. However, this conditioning was not established in rats with cytotoxic lesions made in the RSC one week prior to conditioning. In trace fear-conditioned rats, a number of neurons in the gRSC demonstrated ramping activity, in which the firing frequency gradually increased, synchronous with the onset of the CS tone. This suggests that the gRSC was engaged in interval timing function. Interestingly, this ramping activity was specifically induced by the CS tone, and less ramping activity was observed when a non-CS tone was presented. Furthermore, in rats that had not undergone trace fear conditioning, there were no neurons observed in the gRSC that showed the ramping activity. These results indicate that gRSC neurons demonstrate a plastic change in order to measure various time intervals by utilizing incoming sensory inputs combined with past memory information. The present study suggests a novel function of the gRSC, contributing to interval timing processing.

## 2. Results

### 2.1. Neurotoxic lesions of the gRSC disrupt trace fear conditioning

To examine whether rats can memorize a time interval ranging from the order of seconds to minutes, we employed the trace fear conditioning paradigm in which an US (foot shock) was given after, and temporally separated from a CS (sinusoidal tone, Fig. 1). First, we performed the conditioning using intact rats. The average freezing ratio before, during and after the CS tone presentation was calculated using the behavioral data derived from 11 trace fear-conditioned rats. In the group of intact conditioned rats, the freezing ratios before and during CS tone presentation (31.5 ± 8.57% and 29.9 ± 8.38%, mean ± SEM, respectively) were not significantly different from each other (Fig. 2, P > 0.05, one-way ANOVA with post hoc Bonferroni’s multiple comparison test). The freezing ratio increased immediately after the CS presentation was terminated, and the average value exceeded 60% (63.0 ± 7.52%) which was significantly higher compared to the ratio before and during CS presentation (Fig. 2, P < 0.05, one-way ANOVA with post hoc Bonferroni’s multiple comparison test). This result suggests that rats can acquire the trace fear memory and therefore can memorize the trace time interval. Next, we tested whether neurotoxic lesions in the gRSC produced before the conditioning session impaired the process for acquiring the trace fear memory. In the lesioned rats that were trace fear-conditioned one week after the surgery, the freezing ratio after termination of the CS tone (34.2 ± 11.6%, n=7) showed no significant increase compared to the ratios before and during CS presentation (24.1 ± 10.3% and 18.6 ± 8.92%, respectively, Fig. 2, P > 0.05, one-way ANOVA with post hoc Bonferroni’s multiple comparison test). Compared with the intact rats, the average freezing ratio before and during the CS tone presentation was smaller but not significantly different (P > 0.05, unpaired t-test). On the other hand, the average freezing ratio after the CS tone presentation in the lesioned rats was significantly smaller than that in the intact rats (P < 0.05, unpaired t-test). Thus, the lesioned rats failed to express the freezing behavior which was prominent in the intact rats. These results suggest that the gRSC is necessary for acquiring the trace fear memory.

**Figure 1.**
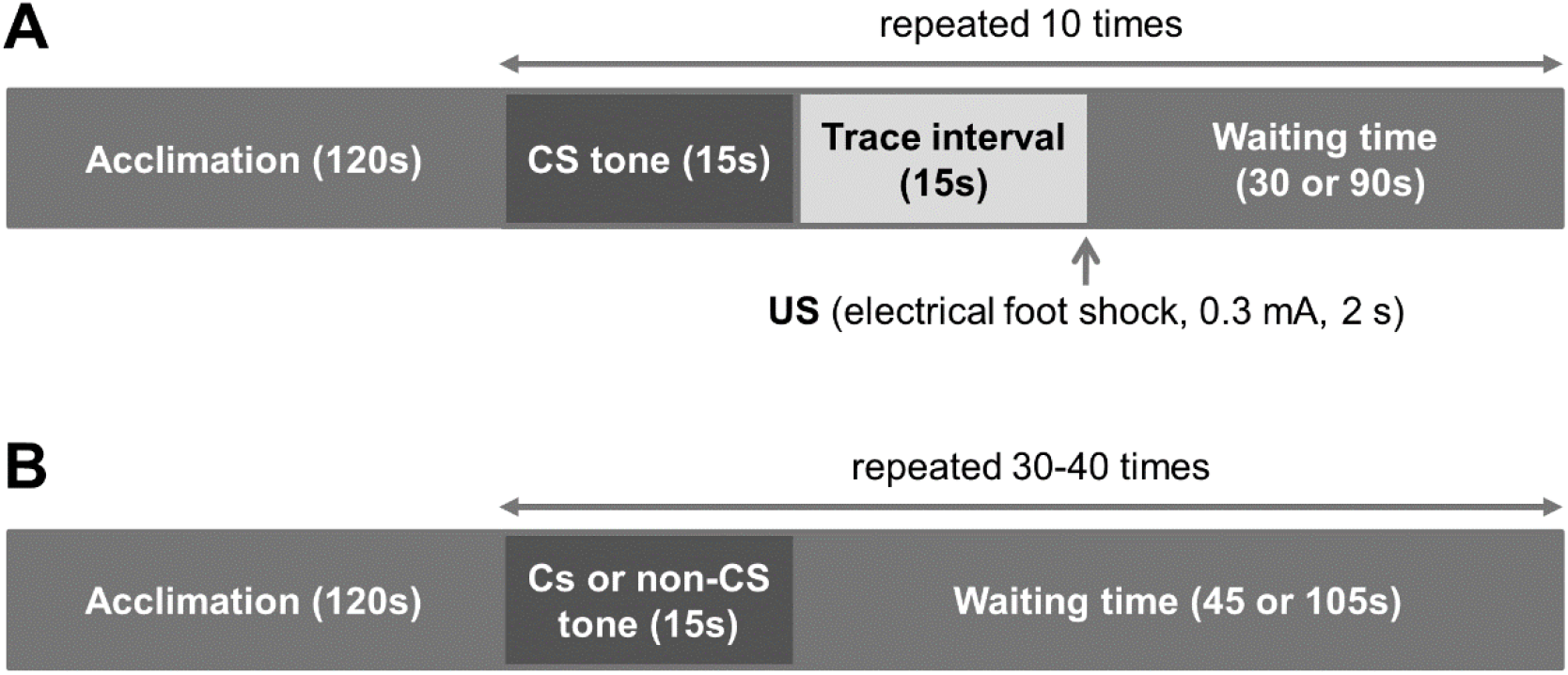
Trace fear conditioning was conducted in a different ‘conditioning box’ placed in a different visual environment from a ‘test box’. A: The conditioning consisted of 10 trials in which a 15-s CS tone was followed by a 15-s stimulus-free trace interval and then a mild foot shock as unconditioned stimulus (US, 0.3 mA, duration 2 s). A waiting time of 30- or 90-s followed the trace interval. B: One week later, rats were tested for freezing behavior in response to the tone presentation (CS or non-CS) in the ‘test box’ with a different visual environment from the conditioning box.

**Figure 2.**
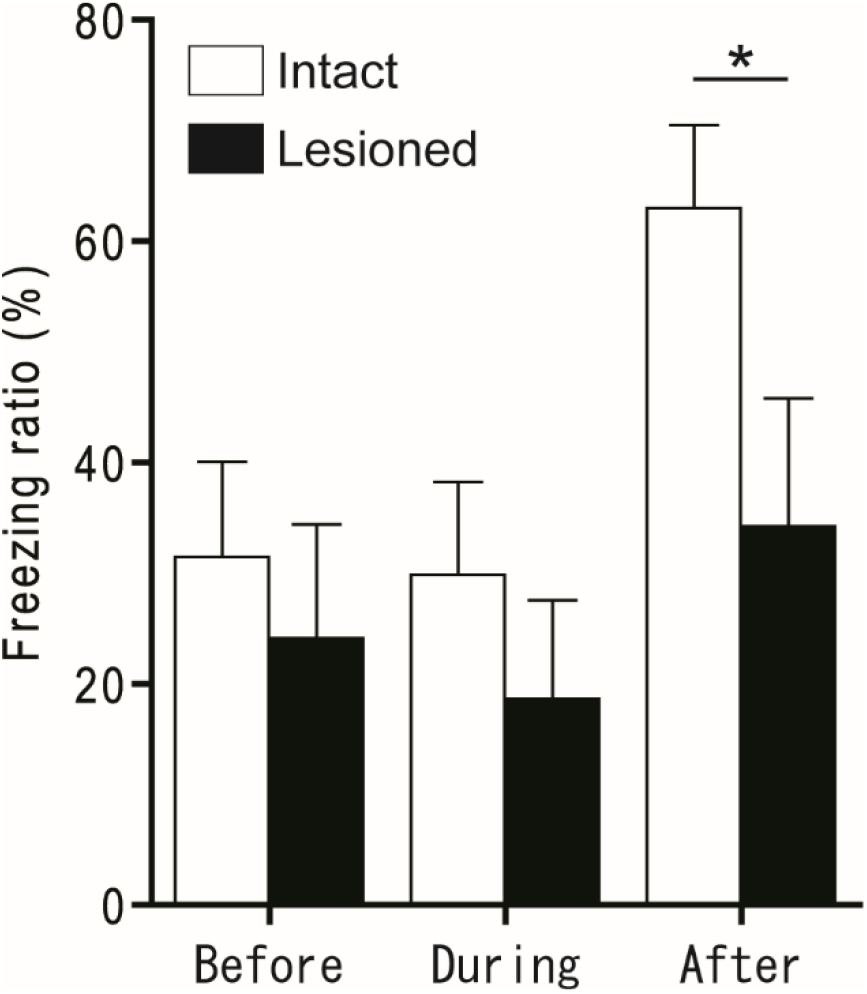
The average freezing ratios obtained from 11 intact and 7 lesioned rats before, during and after the CS tone presentation in the test sessions performed one week after the conditioning. Error bars indicate standard errors of the mean (SEM). The freezing ratio was significantly increased in the intact rats (white boxes) during the time period after the CS tone termination, while no such increase was observed in the lesioned rats (black boxes, see details in the text). The average freezing ratio after the CS for the lesioned rats was significantly smaller than that for the intact rats (asterisk, P < 0.05, unpaired t-test).

### 2.2. Chronic recordings of neural activity in the gRSC

To examine our hypothesis that neural activity in the gRSC encodes interval timing information, we implanted a laboratory-made microdrive electrode array to record activity from the gRSC neurons of freely behaving rats. Recordings were made during a test session, performed one week after trace fear conditioning. Figure 3A indicates representative examples of the spiking activity in gRSC neurons chronically recorded from a free behaving rat and post-stimulus time histograms (PSTH), time-aligned at the beginning of the CS tone stimuli. We considered a change in neural activity to be significant if the deviation of the activity increased or decreased beyond a 95% confidence interval around the overall mean frequency (Fig. 3, grey area in each PSTH). According to this criterion, the spike activity significantly increased in response to presentation of the CS tone (indicated by magenta lines in the figures), and then tended to fall below the average activity level (indicated as dashed lines in the figures). After termination of the CS tone, the activity continuously increased beyond the average level until the next CS tone initiation (Fig. 3A). The existence of such ramping activity in the gRSC strongly indicates that this structure is engaged in temporal processing. We performed chronic recordings in 5 rats. We recorded from 55 gRSC neurons in total and observed ramping activity in 39 of them, with peaks at various time points. However, due to technical problems such as noise signals caused by unexpected body movements of the animals, the recordings were rather unstable, which made further analysis difficult. Therefore, we decided to perform acute multiunit recordings from the gRSC of anesthetized rats to obtain stable recordings and to perform further analyses with a larger sample size.

**Figure 3.**
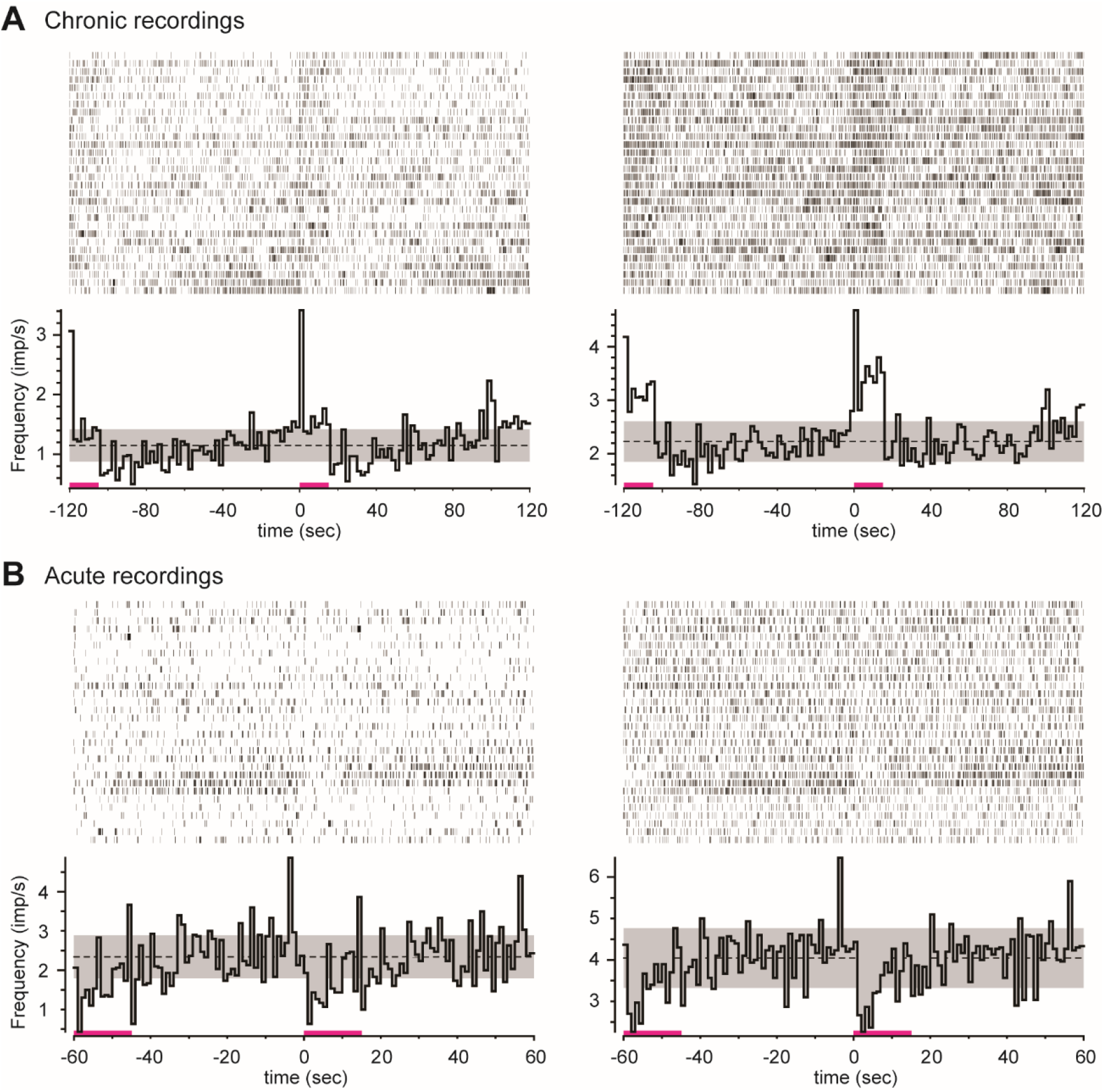
Representative examples of the neural activity recorded from free-moving (A: chronic) and urethane-anesthetized (B: acute) rat gRSCs. Recordings were performed one week after trace fear conditioning. The spiking data from the succeeding 30 test trials aligned (the first trial at the top) with the onset of the CSs (15s tone, magenta bars, t=0, −60 or −120s) are shown in the upper portion of each graph. The lower portion of each graph indicates a post stimulus time histogram (PSTH) for the same 30 trials, in which the mean spike frequency for each time bin (2s for chronic and 1s for acute recordings) was calculated. Magenta bars represent the period of CS tone presentation. Dashed lines indicate the overall mean frequency throughout a test session, and upper and lower limits of the grey area represent 95% confidence intervals.

### 2.3. Acute recordings of neural activity in the gRSC

One week after trace fear conditioning, acute recordings were performed in urethane-anesthetized rats using a 32-channel Neuronexus probe. We could identify 366 neurons from 4 rats that demonstrated various patterns of activity. Among those, we found a considerable number of neurons demonstrating ramping activity synchronized with the onset of the CS tone (Fig. 3B), as was observed in the chronic recordings (Fig. 3A). Of particular interest, we observed a group of neurons that increased their activity until roughly the expected time of the electrical shock, even though only the CS tones were presented and no electrical shocks were applied in the test session (Fig. 4A, left panel). Moreover, there was a strong tendency for the spike activity to be specific to the CS tone. In the same neuron, a non-CS tone (with a different frequency) often failed to induce such discriminative activity (Fig 4A, right panel, see below). We also observed another type of periodic neural activity with the highest peak at various time points from the onset of CS presentation (Fig 4B-D). This activity also tended to be induced specifically by the CS tone (cf. right and left panels in Figs. 4B-D). Since we set 60 seconds as the CS-CS time interval in the acute recordings, we divided the interval into three 20-sec time windows from the onset of CS presentation to further classify the neurons based on when they showed peak responses. Then, we divided the recorded neurons into five groups based on the time at which the largest response was obtained from the onset of the CS tone presentation: namely, neurons that showed the highest activity in 0-20s, 20-40s or 40-60s time windows (Fig. 4A-C), respectively. In addition, a fourth group included neurons that showed peak activity in two or more time windows (Fig. 4D). Lastly, a fifth group included those neurons that showed no significant peak in any time window (Fig. 4E). Table 1 summarizes the numbers and percentages of neurons classified into these 5 categories acutely recorded the from trace fear-conditioned and naïve rats. In rats subjected to trace fear conditioning, when the CS tone was presented, 15% of neurons (55/366) showed peak activity within the 20-40s time window (the electrical shock was expected to arrive at 30s). On the other hand, when the non-CS tone instead was presented to the same conditioned rats, this percentage reduced to 4.4% (16/366) in the same neuronal population. In the 0-20s time window, the non-CS tone induced peak activity in 25% of neurons (92/366) whereas the CS tone only induced peak activity in 9% of neurons (33/3666). We performed similar acute recordings using two “naïve” (non-conditioned) rats and recorded stable neural activity from 191 neurons. We found that 28% (53/191) of the neurons showed peak activity in the 0-20s time window induced by the tone presentation, comparable with that observed in the conditioned rats with the non-CS tone (25%). These findings suggest that about 25-30% of gRSC neurons in rats respond to non-CS tones, regardless of trace fear conditioning experience. In addition, in naïve animals, very few neurons showed the largest activity in the 20-40s (0.52%, 1/191) or 40-60s (0%, 0/191) time windows and more than 60% (116/191) of neurons demonstrated no significant increase in neural activity in response to the tone presentation (< >95% of the confidence interval). A Chi-square test showed that the distribution patterns of the number of cells in the five categories for CS, non-CS and naïve groups were significantly different from one another (Chi-square; 158.9, df; 8, P < 0.0001, Table 1). Our observations strongly suggest that the changes in neural activity observed during the test session after the establishment of trace fear conditioning was specific to the CS tone. In addition, the CS tone presentation induced peak activity in the 20-40s time window, when the noxious stimulus was expected. These results indicate that the trace fear conditioning invoked plastic changes in neural activity of the rat gRSC so that a significant number of neurons showed an increase their activity around the time when the electrical foot shocks were expected to arrive.

**Figure 4.**
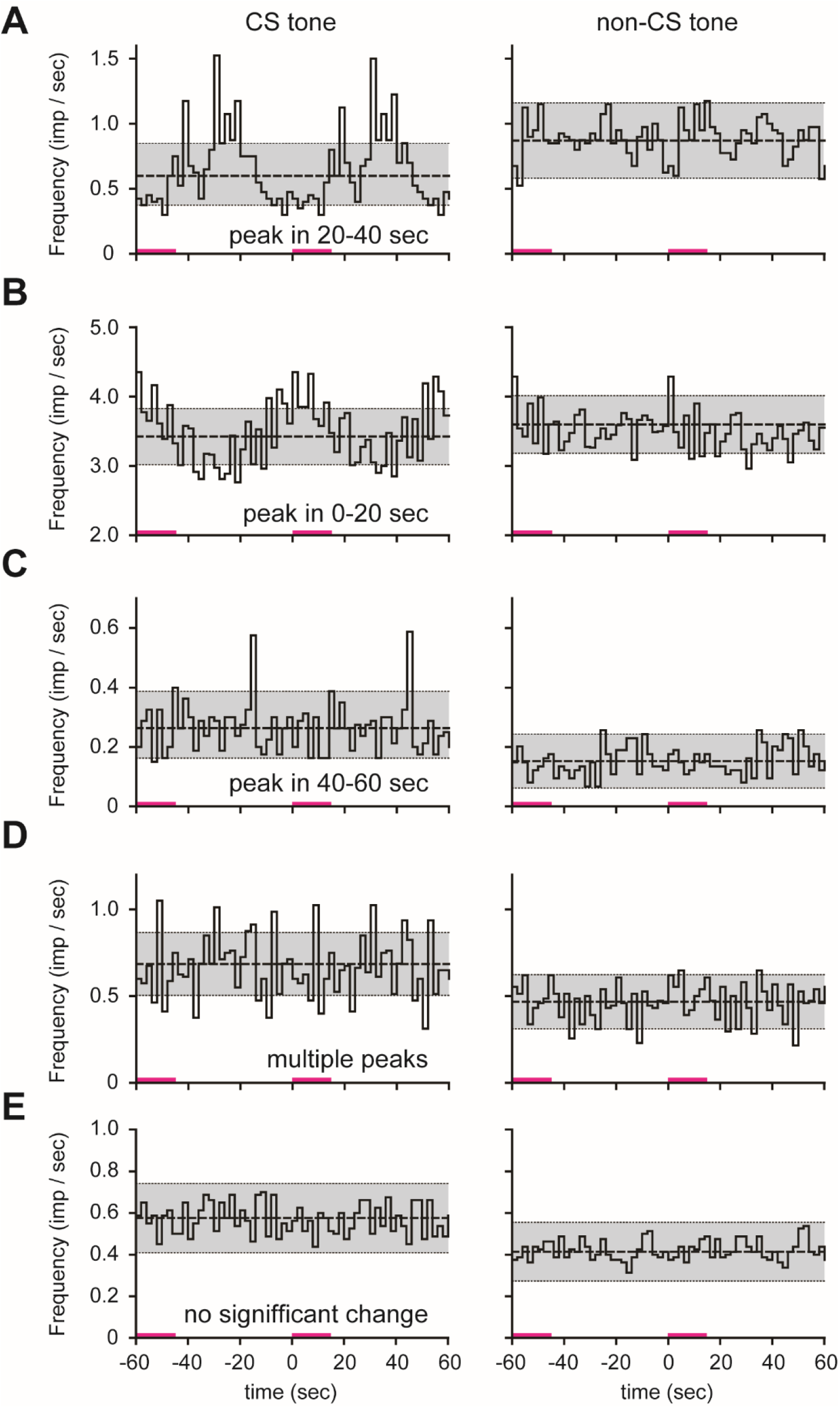
PSTH examples of the gRSC neurons acutely recorded from urethane-anesthetized rats. Left and right panels in each row represent the PSTHs calculated from the neural activity (30 trials, time bin, 2s) recorded from the same neuron in the gRSC, in response to CS and non-CS tone stimuli, respectively. Magenta bars indicate the period for tone presentation. Dashed lines indicate the overall mean frequency, and upper and lower limits of the grey area represent 95% confidence intervals. A: The CS tone tended to induce peak activity during the 20-40 sec interval in a larger number of cells than the non-CS tone. B and C: Likewise, a significant number of neurons demonstrated the peak activity during 0-20s or 40-60s time windows. As observed in the chronic recordings (Fig. 3A), similar ramping activity demonstrating the highest peak activity in different time points from the onset of the CS tone was observed in many neurons in the acute recordings (A-C). D: Multiple peak activity (oscillation) was also invoked by CS tones, while non-CS tones did not induce such oscillatory responses. E: In some neurons, neither CS nor non-CS tones induced significant ramping activity (see Table 1 and Fig. 5).

**Figure 5.**
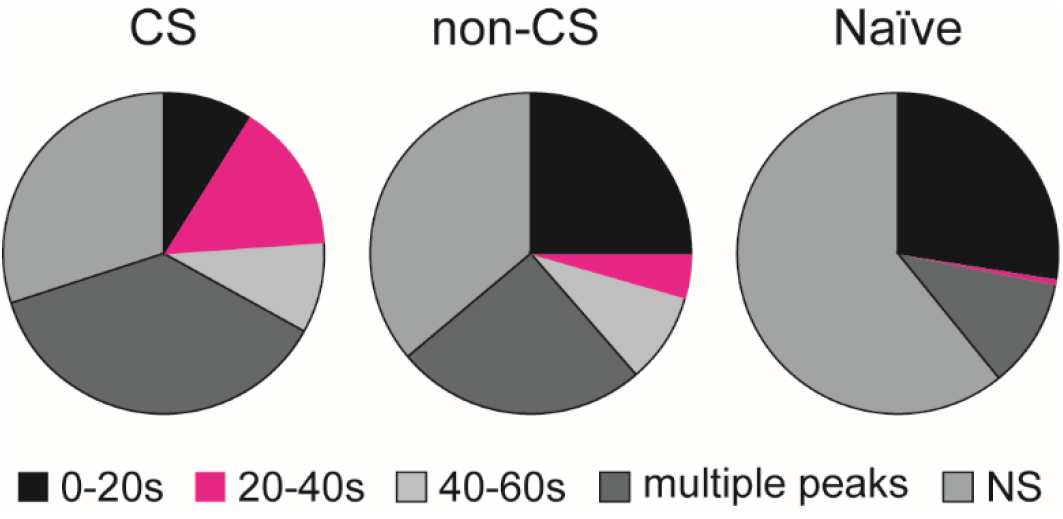
The percentage breakdown of neurons showing peak activity in different time windows recorded from trace fear-conditioned rats and naïve rats (see Table 1). Note that in conditioned rats, the percentage of neurons demonstrating peak activity during the 20-40s time window (when the US foot shock was expected to arrive; shown in magenta) was much larger than that observed in the non-CS or naïve groups.

**Table 1.** Number and percentage of cells categorized in 5 activity patterns. The figures on the table indicate the number of the cells classified in 5 groups according to the location of the highest peak measured from the onset of stimulus tone, as indicated on the top row. The figures in parentheses indicate the percentage of the cells to the total number of cells recorded (366 for CS and non-CS groups and 191 for the naïve group).

|  | 0-20 s | 20-40s | 40-60s | Multiple<br>peaks | No significant<br>peaks |
| --- | --- | --- | --- | --- | --- |
| CS | 33 (9.0%) | 55 | 33 | 136 | 109 (29.8%) |
| Non-CS | 92 | (15.0%) | (9.0%) | (37.2%) | 132 (36.1%) |
| Naïve | (25.1%) | 16 (4.4%) | 34 | 92 (25.1%) | 116 (60.7%) |
|  | 53 | 1 (0.5%) | (9.3%) | 21 (11.0%) |  |
|  | (27.7%) |  | 0 (0%) |  |  |

## 3. Discussion

### 3.1. Possible gRSC function in interval timing during trace fear conditioning

In the present study, we performed trace fear conditioning using rats to test the possibility that the RSC encodes time intervals on the order of seconds to minutes, and found the following results. First, rats subjected to the trace fear conditioning learned the time intervals between the CS and CS or CS and US, and began to express a freezing response as an adaptive behavior. However, this associative learning was not established in rats with cytotoxic lesions in the RSC one week prior to conditioning. Thus, the RSC is necessary for the acquisition of the trace fear memory. Second, in chronic and acute recording experiments, many neurons in the gRSC of trace fear-conditioned rats demonstrated a progressive increase in neuronal firing frequency; the ramping activity was synchronized with the onset of the CS tone stimuli. Third, in acute recording experiments, the gRSC of trace fear-conditioned rats contained a distinct neural population that was most active at the time when the electric foot shocks were expected to arrive. This neuronal activity was specific to the CS tone and was not responsive to the non-CS tone. Furthermore, in the gRSC of naïve rats that had not undergone trace fear conditioning, few neurons exhibiting such activity were observed. These results suggest that the rat gRSC is involved in interval timing mechanisms that memorize and recall time lengths on the order of seconds to minutes, which is necessary for carrying out appropriate adaptive behaviors.

We have previously revealed that small pyramidal neurons in the superficial layer of the rat gRSC are late-spiking (LS) neurons that generate action potentials with a time delay in response to inputs (Kurotani et al., 2013). Given this LS firing property, LS neuron alone can function as a time integrator on the order of seconds. It may also be possible to encode much longer time lengths by cascading neighboring LS neuron modules. It has been shown by a neural model simulation that neural circuits containing LS neurons can encode time on the order of seconds (McGann and Brown, 2000). Morphological analyses have shown that LS neurons extend axonal lateral branches hundreds of microns along the superficial layers and may form synaptic connections to nearby LS neurons (Ichinohe et al., 2003, Kurotani et al., 2013). Using optical imaging of neural activity and current source-density analysis, we showed that electrical stimulation in layer 1a, mimicking input from the anterior nucleus of the thalamus, induced neural excitation in the superficial layer of the gRSC. Then, this excitation propagated transversely within the superficial layers (Nixima et al., 2013, 2017). It has been reported in previous studies using a double whole-cell recording technique that LS neurons did not form connections with one other (Brennan et al., 2020, Robles et al., 2020). These findings are inconsistent with our observations described above. This discrepancy may be due to the fact that in double-patch recordings, it is possible to record only from two neurons with close proximity, and it is difficult to record from cells that are far apart.

Figure 6 shows a hypothetical schematic diagram of the internal circuit of the gRSC that buffers the time delay, according to the previous findings and the present results. LS neurons in the superficial layer of the RSC receive a variety of sensory inputs from the thalamus within layer 1 (Sripanidkulchai and Wyss, 1986, Wyss et al., 1990, van Groen and Wyss, 1990, Shibata, 1993, Ichinohe et al., 2003, Odagiri et al., 2011, Sugar et al., 2011). In the case of trace fear conditioning, because of the time difference between the CS tone and US foot shock, a time delay circuit is required for associative learning between the CS and US to be established. It has been suggested that LS cells forming patchy clusters in the superficial layers of the gRSC are synaptically connected with one another, and this cascading network can generate a time delay on the order of seconds (Ichinohe et al., 2003, Kurotani et al., 2013, Nixima et al, 2013, 2017). It was also shown that the excitation induced by layer 1a stimulation propagated vertically to the deeper layers (Nixima et al., 2017). We propose that in trace fear conditioning, information from the presented CS tone propagates horizontally through the superficial layer with a time delay caused by LS cells (Fig.6, thick horizontal black arrow). After a certain time delay, the US foot shock is applied, and information from both the CS and US converge at a single site where Hebbian synaptic plasticity takes place (“Hebbian learning” in Fig. 6). After the association is established, signal transmission at this site is enhanced, and information from the CS is more easily transmitted to the deep layers below than to other sites. Pyramidal neurons in deep layers receive inputs from the dorsal hippocampus (Wyss and van Groen, 1992, van Groen and Wyss, 2003, Sugar et al., 2011, Yamawaki et al., 2019a and 2019b). Finch et al. (1984) reported that thalamic and hippocampal inputs were integrated in layer 5 pyramidal neurons in the gRSC. There is much evidence demonstrating the involvement of the hippocampus in trace fear conditioning (D’Adamo et al., 2004, for review, Raybuck et al., 2013). During a test session in trace fear conditioning, a recalled fear memory is thought to be transmitted to the pyramidal neurons via the hippocampal-retrosplenial pathway (“HFC input” in Figure 6). The delayed CS tone information and the recalled fear memory thus converge at the site where Hebbian learning occurs, which induces the enhancement of synaptic transmission. As a result, freezing behavior is expressed with an appropriate time delay. The circuit shown in Fig. 6 may be oversimplified, and indeed there are many subtypes of inhibitory interneurons in the gRSC playing crucial roles in modifying the activities of those excitatory neurons (Nixima et al, 2017, Brenann et al, 2020). However, our observations at least seem to support the basic concept illustrated in Fig. 6 that a time-delay circuit is somehow formed in the rat gRSC.

**Figure 6.**
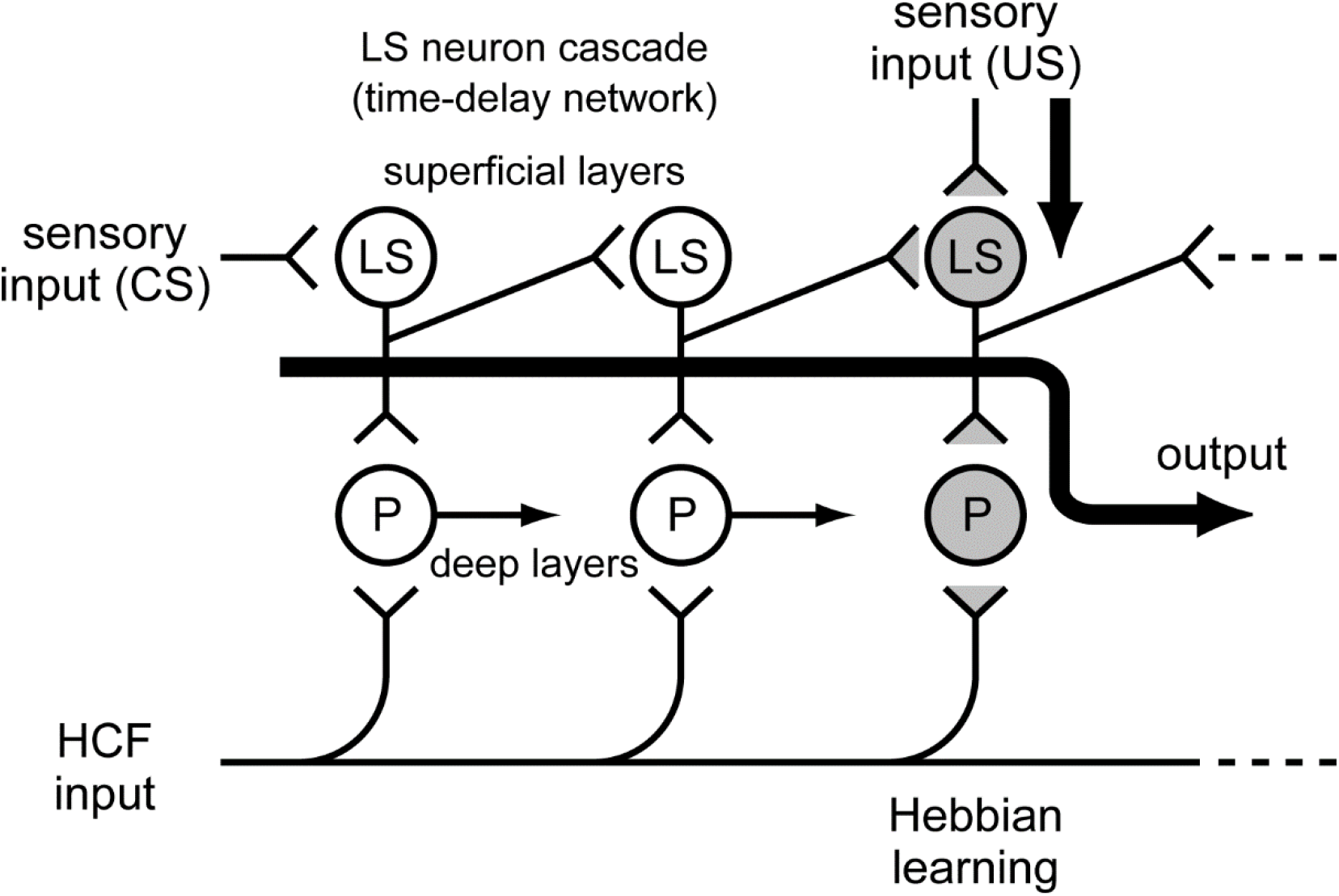
Schematic drawing of the time-delay mechanism embedded in the local circuit of the rat gRSC. Various sensory inputs converge on the late-spiking neurons (LS) in the superficial layers of the gRSC. LS neuron cascades form horizontal connections in the superficial layers, and this network can generate time delays of the order of seconds due to their late-spiking firing properties. In the case of trace fear conditioning, there would be a certain place in this network, where information from the delayed tone stimulus (CS) and the foot shock signal (US) come together at the same time. Then, only there does the Hebb-type synaptic strengthening take place (gray synapses). The strengthened synapses transmit the CS signal with a time delay to the pyramidal neurons (P) in the deep layers just below the LS neuron module. During the test session, recalled memories of the fear are transmitted from the hippocampus (HCF input), and further Hebbian synaptic reinforcement occurs in the deep layer pyramidal cells. This results in the expression of a freezing behavior with an appropriate time delay.

### 3.2. Known Features of the RSC and interval timing

RSC functions are diverse and are deeply involved in spatial learning and memory, and navigation (Aggleton et al. 2010; Chen et al. 1994; Cooper and Mizumori 2001; Dumont et al. 2010; Garden et al. 2009; Pothuizen et al. 2008; Vann et al. 2009; Vogt and Miller 1983), as well as in episodic memory acquisition and recall (Aggleton and Pearce, 2001, Vann et al., 2009, Robinson et al, 2014). Imaging studies in humans show that RSC is activated by recalling autobiographical memory (Vann et al., 2009), scene perception (MacEvoy and Epstein 2007), mapping tasks, and imagining future selves (Schacter et al., 2007). It has also been reported that navigational and episodic memory is impaired in humans after RSC injury (Maeshima et al., 2001, Ino et al., 2007, Valenstein et al., 1987). The link between spatial navigation and episodic memory has been debated in recent years (Takahashi 2018). Episodic memory is characterized by the fact that it contains not only information about the place where the event occurred, but also the order in which things occurred, namely, chronological information. It has also been suggested that visuospatial navigation is closely related to other forms of memory processing, including contextual processing and episodic memory, as it contains information on the location of objects that appear sequentially along the path of travel and the temporal changes of environmental factors (Eichenbaum et al., 1999). That is, the RSC play an important role in integrating various stimuli from the surrounding physical environment in a spatiotemporal context (Morris, 2001). Therefore, it is plausible to assume that the RSC has an embedded system that processes not only spatial information but also temporal information.

The lesion experiment in the present study showed that the RSC was necessary for the acquisition of the trace fear memory. It remains unclear as to whether it is also sufficient for acquisition, or whether there are other factors involved. The time interval between the CS and US used in the present study was 15 s, and the interval between the CS and CS was 60 to 120 s, which are all in the range of times processed by interval timing mechanisms. Several previous studies showed that the RSC contributes to interval timing (Todd et al., 2015, Hayashi et al., 2020). In humans, there have been case reports that patients with RSC damage are unable to judge the order in which events occurred (Bowers, et al, 1988). Considering these facts together with the time-delayed firing properties of LS neurons and the macroscopic information transfer properties of local circuits in the RSC (Nixima et al, 2013, 2017), it is plausible that the RSC has the ability to encode time lengths on the order of seconds to minutes, which may contribute to episodic memory acquisition, recall, and visuospatial navigation.

Temporal processing for episodic memory in the hippocampus occurs at the population level and is discrete (Manns et al., 2007, Pastalkova, 2008, MacDonald et al., 2011). In contrast, we found that interval timing processing in the gRSC is achieved at the single-neuron level and is temporally continuous (displayed as ramping activity). The RSC has a mechanism for transferring memory engrams containing episodic memories from the hippocampus to the neocortex for long-term storage as memories (Lesburgueres et al., 2011, Tanaka et al., 2014, Kitamura et al., 2017). In this process, the discrete temporal information of the hippocampus and the continuous temporal information of the gRSC may be used in a complementary manner. For example, information on the location and order of appearance of specific targets during navigation may use the discrete information system of the hippocampus, while the time required to travel between two targets may be continuously measured by the timing mechanism of the gRSC via ramping activity. Then, the integrated memory would be transferred to the neocortex for storage as a long-term memory (Sousa et al., 2019, Nitzan et al, 2020).

### 3.3. Comparison with previous models of interval time processing

A variety of phenotypes for interval timing have been reported. In hippocampal CA1, there exists a group of pyramidal neurons that are sequentially active during tasks that require animals to memorize delay intervals (MacDonald et al., 2011, 2013). These “time cells” are active at specific time points, and their activity is discrete and represented as firing of a certain population of neurons. A different mechanism has been proposed for timing detection, by the medium spiny neuron (MSN) in the basal ganglia (Mattel et al. 2003, Buhusi and Meck, 2005). It is hypothesized that MSNs centrally receive the outputs of many cortical oscillators and utilize synaptic plasticity to capture the state of these inputs at the moment of an event’s beginning and ending. In addition to these phenotypes, ramping activity is also regarded as a distinct form of time representation in the brain. In the present study, ramping activity was recorded from a single gRSC neuron in rats subjected to trace fear conditioning with CS-CS intervals ranging from 60 to 120 seconds. In addition, ramping activity was also observed in a considerable number of neurons (Table 1). This activity was initiated by the CS tone and showed the highest peak at the time when the US electrical foot shock was expected to arrive (15 s after the termination of the CS tone in our experiments). This activity was specific to the CS tone and was not induced by the non-CS tone (Fig. 4). Furthermore, this activity was only recorded from rats that had been subjected to trace fear conditioning. These results indicate that the gRSC has a neural mechanism to continuously measure the time from the onset of a particular stimulus. A unique feature of the interval timing mechanism in the gRSC is that it can be used both retrospectively and prospectively. In a test session of trace fear conditioning, experientially learned time interval memories were referenced and then behavior (freezing) was expressed. In this case, the interval timing mechanism was used retrospectively. In an operant conditioning experiment in which rats were trained to press a lever for a specific length of time to receive a reward, neurons that fired specifically during the lever pressing behavior were observed in the gRSC of the conditioned rats (Tanaka, Master’s Thesis submitted to the faculty of the Graduate School of Arts and Sciences, The University of Tokyo). This suggests that the gRSC also contributes to prospective time generation.

In summary, we revealed in the present study that the rat gRSC can process time intervals ranging tens to hundreds of seconds by utilizing the time integration properties of LS neurons. Taken together with the results of our previous studies, we have shown the functional significance of the gRSC in interval timing processing, from the cellular level to the behavioral level. However, it is still unclear what synaptic mechanisms are responsible for storing and referencing the time intervals between events that are separated in time, resulting in the induction of appropriate responses and behaviors. Further investigation of this will be required.

## 4. Materials and Methods

### 4.1. Subjects

All experiments and procedures were approved by the Institutional Animal Care and Use Committee of the University of Tokyo. Male Wistar rats, 12-24 weeks old (body weight, 290-320g) were used for the experiments. Rats were individually housed in polycarbonate cages with free access to water and 20 g of food pellets per day, in a 12 h-12 h light-dark cycle.

### 4.2. Trace fear conditioning

Trace fear conditioning was performed using an MK-450 fear conditioning box (Muromachi Kikai Co., Ltd., Tokyo, Japan). The size of the conditioning chamber was 280 mm (width) x 220 mm (depth) x 500 mm (height) with a speaker and a white LED light on the side wall. The chamber equipped 17 stainless steel rods (4mm in diameter, 15 mm apart) at the bottom connected to an electrical shock generator for applying foot shocks. To monitor an animal’s behavior, a digital video camera was placed at the top of the chamber. The video signal was fed to a computer and used to calculate the freezing ratio via video tracking software (CompACT VAS/DV, Muromachi Kikai Co., Ltd.). Freezing was defined as complete immobility of the animal for more than 3 seconds, and the freezing ratio was calculated as the percentage of time spent freezing, starting from the beginning of the test session, and measured in 15-sec time bins. Timing for generation of tones, lights and foot shocks was also controlled by the same tracking software.

During the first five days of the experiment, each rat was transferred to a ‘test chamber’ for 10-20 minutes per day and tones used for the conditioned stimulus (CS) were randomly presented to acclimate the animal to the experimental environment. The test chamber (560 mm (width) x 440 mm (depth) x 1000 mm (height)) was a laboratory-made cage whose size, wall color, and surrounding visual environment differed from those of the conditioning chamber, to minimize the effect of contextual cues. On day 8, each rat was transferred to the conditioning chamber and the trace fear conditioning procedure was conducted. The conditioning consisted of 10 trials in which a 15-s CS tone (2, 8, or 16 kHz sinusoidal wave at 85 dB) was followed by a 15-s stimulus-free trace interval and then a mild foot shock as the US (0.3 mA, duration 2 s, Fig. 1A). The interval of CS tone presentation was fixed at either 60 or 120 sec. After the conditioning session, rats were returned to their home cages. One week later (on day 15), rats were tested for freezing to the CS or non-CS tone in a test chamber that had a different size, color, and surrounding environment from the chamber used in the conditioning session (Fig. 1B).

### 4.3. Surgery

A total of 18 rats (7 for cytotoxic lesions, 5 for chronic and 6 for acute neural recordings) were used for the experiments requiring surgery. Each rat was anesthetized with pentobarbital (40 mg/kg i.p.) and placed on a stereotaxic apparatus (SR-6R-HT, Narishige, Tokyo, Japan). A midline incision was made in the scalp and the skull was exposed. Lidocaine jelly (XYLOCAINE® JELLY 2%, AstraZeneca K.K., Osaka, Japan) was also applied to the incision for local anesthesia. During the surgery, anesthetic level was occasionally tested by tail-pinch method, and maintained by additional administration of 0.5-1% isoflurane. For cytotoxic lesions of the gRSC, small holes were drilled on the skull at 8 different points along the rostrocaudal axis (coordinates measured from bregma: AP −3.6, −4.7, −5.8 and −6.5 mm, ML ±0.6 mm, DV −1.6 to 2.0 mm from the skull). Lesions were made by bilateral injection of 0.2-0.3 μl of phosphate buffered saline containing N-Methyl-D-Aspartate (10mg/ml) with a Hamilton syringe (80300, Hamilton Company, Reno NV) attached to a syringe pump (KDS-310-PLUS, KD Scientific, Holliston MA, flowrate: 0.05 μl/min). One week after the surgery, the same conditioning procedure as done for intact rats was conducted for the lesioned rats. After the completion of the behavioral experiments, rats were deeply anesthetized with pentobarbital (100 mg/kg i.p.), and transcardially perfused with 4% paraformaldehyde (in phosphate buffered saline). The extent of damage caused by NMDA injection was evaluated in Nissl-stained coronal sections (thickness 50 μm), and data obtained from animals with ∼ 70% or greater damage in gRSC were selected for the analysis of the behavioral experiment.

A laboratory-made tetrode array combined with a microdrive was implanted and used for the chronic neural recordings from the gRSC. The fabrication procedure for the electrode assembly was the same as described by Takahashi and Sakurai (2005). Briefly, the tetrode consisted of four tungsten wires (diameter, 15 μm) inserted in a 33-gauge stainless steel guide tubing, and then either 2 or 4 tetrodes were attached to a microdrive at 0.5-0.8 mm intervals. Same as the lesion surgery, each rat was anesthetized and placed in the stereotaxic apparatus. A square window (AP −4 to −7.5 mm, ML 0.5 to 2.5 mm) was drilled in the right side of the skull. After incising the dura mater with a 27-gauge syringe needle, the tetrode array was positioned so that the electrode tip touched the cortical surface 1 to 1.2 mm lateral to the midline. Because the gRSC lies bilaterally on the medial surface of the cerebral hemispheres and the interhemispheric fissure is covered by the superior sagittal sinus, it was difficult to insert the electrodes perpendicular to the gRSC. Therefore, the electrode was tilted 30 degrees lateral from the median plane. The antero-posterior position of the electrode tips was about 5 to 6.5 mm posterior to bregma. The window in the skull was filled with sterilized white petrolatum to prevent acrylic cement from directly touching the dura matter, and then the whole tip of the electrode assembly was embedded in acrylic cement. The rats recovered for 7 to 10 days after the surgery, and then the trace fear conditioning process was initiated. One week after conditioning, chronic neural recordings were conducted in freely moving rats during the test session.

For acute recordings, each rat was anesthetized with urethane (1 to 1.5 g/kg i.p.) and placed in the stereotaxic apparatus. A midline incision was made in the scalp and the skull was exposed. Lidocaine jelly was also applied to the incision for local anesthesia. A square window (AP −4 to −7.5 mm, ML 0.5 to 2.5 mm) was drilled in the right skull and a small incision was made in the dura matter, approximately 1 to 1.5 mm lateral to the midline. After this stereotaxic surgical procedure, a small stainless steel screw was installed at the anterior part of the skull, and a stainless steel head post was fixed to the screw by acrylic cement. Then the ear bars were removed so that the sound stimulation was readily audible to the rat. A 32-channel Neuronexus probe (Neuronexus Technologies, Inc. Ann Arbor, MI) was inserted into the cortex with an angle of 30 degrees lateral from the medial plane as was the case in the chronic recordings. The antero-posterior position of the electrode tips was about 5 to 6.5 mm posterior to the bregma. The electrode array was mounted on a water-driven micromanipulator (WM-105, Narishige, Tokyo, Japan) and was slowly advanced about 2 mm to access gRSC neurons.

### 4.4. Recording neural activity in the gRSC

Both in the chronic and acute experiments, multiunit neural activity was recorded using a Plexon Recorder system (Plexon Inc., Dallas, TX). Offline spike sorting was performed using a valley-seek and k-means clustering algorithm (Offline Sorter; Plexon), followed by manual confirmation of sorting validity. Peri-event histograms were constructed for each unit activity by Neuroexplorer (Nex Technologies, Madison, AL). The spiking data from the succeeding 30-40 test trials was aligned with the onset of the CS of each trial and the mean spike frequency for each time bin (1-2s) was calculated.

## Declaration

The authors declare no conflicts of interest associated with this manuscript.

## Author contributions

**Tohru Kurotani:** Conceptualization, Methodology, Investigation, Data curation, Formal analysis, Writing - original draft, Writing - review & editing, Funding acquisition.

**Ken’ichi Nxima:** Conceptualization, Methodology, Investigation.

**Tomohiro Tanaka:** Investigation.

**Yoshio Sakurai:** Methodology, Resources.

**Kazuo Okanoya:** Conceptualization, Supervision, Writing - review & editing, Funding acquisition.

## Acknowledgements

We thank Ms. Keiko Kikuchi for technical support.

## Funding

This work was supported by the Okanoya Emotional Information Project of the Japan Science and Technology Agency (JST-ERATO), RIKEN Brain Science Institute (BSI), KAKENHI no. 26119509 to K. Okanoya, and KAKENHI no. 22500370 to T. Kurotani.

## Notes

### Competing Interest Statement

The authors have declared no competing interest.

